# MyD88-dependent TLR signaling oppositely regulates hematopoietic progenitor and stem cell formation in the embryo

**DOI:** 10.1101/2021.07.21.453199

**Authors:** Laura F. Bennett, Melanie D. Mumau, Yan Li, Nancy A. Speck

**Author notes:** Correspondence: Nancy A. Speck, Abramson Family Cancer Research Institute and Department of Cell and Developmental Biology, Perelman School of Medicine, University of Pennsylvania, Philadelphia, PA 19104, USA;. These authors contributed equally to this work (Laura F. Bennett, Melanie D. Mumau).

## Abstract

Hemogenic endothelial (HE) cells in the dorsal aorta undergo an endothelial to hematopoietic transition (EHT) to form lympho-myeloid biased progenitors (LMPs), pre-hematopoietic stem cells (pre-HSCs) and adult-repopulating HSCs. These briefly accumulate in intra-arterial hematopoietic clusters (IAHCs) before being released into the circulation. It is generally assumed that the number of IAHC cells correlates with the number of HSCs. Here we show that changes in the number of IAHC cells, LMPs, and HSCs can be uncoupled. Mutations impairing MyD88-dependent toll-like receptor (TLR) signaling decreased the number of IAHC cells and LMPs but increased the number of HSCs in the aorta-gonad-mesonephros region of mouse embryos. TLR4 -deficient embryos generated normal numbers of HE cells but the proliferation of IAHC cells was decreased. Loss of MyD88-dependent TLR signaling in innate immune myeloid cells had no effect on IAHC cell numbers. Instead, TLR4 deletion in endothelial cells recapitulated the phenotype observed with germline deletion, demonstrating that MyD88-dependent TLR signaling in endothelial cells and/or in IAHCs regulates the balance between generating LMPs and HSCs.

**Summary Statement:** Toll-like receptor signaling in endothelial cells restricts the number of hematopoietic stem cells but increases the number of committed progenitors and intra-arterial hematopoietic cluster cells by promoting their proliferation.

## Introduction

Hematopoietic ontogeny occurs in multiple waves in which hematopoietic stem and progenitor cells (HSPCs) with different potentials differentiate from hemogenic endothelial (HE) cells via an endothelial to hematopoietic transition (EHT). HE cells in the extra-embryonic yolk sac undergo EHT to become committed erythro-myeloid (EMP) and lymphoid progenitors, while HE cells in the caudal arteries (dorsal aorta in zebrafish; dorsal aorta, vitelline and umbilical in mouse) differentiate into multipotent, lympho-myeloid biased progenitors (LMPs), HSCs, and HSC precursors called pre-HSCs (Auerbach et al., 1996; Hadland and Yoshimoto, 2018). Following EHT, the newly formed HSPCs in the major arteries briefly accumulate in intra-arterial hematopoietic clusters (IAHCs) consisting of cells that uniformly express endothelial markers, hematopoietic transcription factors such as RUNX1, and high levels of Kit (Bollerot et al., 2005). At embryonic day (E) 10.5, when IAHC numbers peak, the IAHC cells contain several hundred LMPs, approximately 25 pre-HSCs, and ∼0.03 adult-repopulating HSCs (Kumaravelu et al., 2002; Li et al., 2014; Rybtsov et al., 2016). At E11.5, IAHCs contain approximately 65 pre-HSCs and 1 adult repopulating HSC (Kumaravelu et al., 2002; Rybtsov et al., 2016). Hence the IAHCs are heterogeneous, containing HSPCs with unique differentiation and engraftment potentials, and the balance between the different types of HSPCs changes during development.

Innate immune and inflammatory signaling pathways have been shown to regulate the number of HSPCs generated in the arteries of zebrafish and mouse embryos. Mutations in or morpholino knockdowns of genes encoding type I and II interferons (Li et al., 2014; Sawamiphak et al., 2014), tumor necrosis factor (TNF) (Espin-Palazon et al., 2014), inflammasomes (Frame et al., 2020), interleukin 1β (IL-1β) (Frame et al., 2020), granulocyte colony stimulating factor (G-CSF) (He et al., 2015), toll-like receptor 4 (TLR4) (He et al., 2015), nuclear factor kappa B (NF-kB) (Espin-Palazon et al., 2014; He et al., 2015) and RIG-1-like receptors (Lefkopoulos et al., 2020) reduced the number of myb:GFP^+^ kdrl:mCherry^+^ HSPCs in transgenic reporter zebrafish embryos and decreased the levels of *runx1* and *myb* transcripts in the dorsal aorta (DA). Innate immune signaling is initiated by binding of ligands to pattern recognition receptors (PRRs). PRRs recognize pathogen-associated molecular patterns (PAMPs), which are small molecules common to a particular class of infectious agents, and host molecules referred to as danger-associated molecular patterns (DAMPs) which are endogenous proteins or small molecules that are released from the cell or the extracellular matrix in response to cell damage, stress, or death. Although, in theory, PAMPs from infectious agents might be responsible for activating innate immune signaling in mammalian or pre-hatched zebrafish embryos, the most plausible ligands for PRRs in the relatively sterile environment of the mammalian embryo are DAMPs.

Activation of toll-like receptors (TLRs) drives the production of type I interferons, TNF, IL-1β and G-CSF, and therefore directly or indirectly stimulates multiple inflammatory pathways (Monlish et al., 2016). TLRs signal through two intracellular pathways that involve either the MyD88 (myeloid differentiation primary response protein) and TIRAP (toll-interleukin 1 receptor domain containing adaptor protein) or the TRIF (toll IL-1 receptor (TIR)-domain-containing adaptor inducing interferon b) and TRAM (translocating chain-associated membrane protein) adaptor proteins. Signaling from TLR4 on the plasma membrane through MyD88/TIRAP results in the nuclear localization of the CREB, AP1, and NF-κB transcription factors and expression of inflammatory cytokine genes such as TNF, IL-1β and G-CSF. Signaling through endosomal TLR4, TRIF/TRAM and the interferon regulatory factors (IRFs) primarily activates the expression of interferon genes. Morpholino knockdown of the genes encoding TLR4 (*tlr4bb*) and MyD88 (*myd88*) in zebrafish embryos reduced the expression of *runx1*, the number of myb:GFP^+^ kdrl:mCherry^+^ HSPCs in the DA, and HSPCs in secondary sites of colonization including the caudal hematopoietic tissue and thymus (He et al., 2015). However, markers that discriminate between committed progenitors (LMPs) and pre-HSCs/HSCs are not available for zebrafish embryos, therefore how TLR4 signaling affects the balance between them is difficult. Understanding how this balance is regulated is important for specifically optimizing generation of the desired hematopoietic stem or progenitor cells from arterial HE cells *ex vivo*.

Here we examine TLR signaling to determine in which cells it is required, through which intracellular adaptors it signals to regulate arterial HSPC ontogeny, and the types of HSPCs it regulates. We show that signaling through the MyD88 arm of the TLR pathway increases the number of IAHC cells and LMPs, while restricting the number of multi-lineage adult-repopulating HSCs.

## Results and Discussion

### TLR4 signaling regulates RUNX1 levels and the number of IAHC cells

It was reported that loss of TLR4 reduced expression of *runx1* in the DA of zebrafish embryos and the number of RUNX1^+^ cells in the DA of mouse embryos (He et al., 2015). To examine whether the reduction in *runx1* expression could be due to fewer RUNX1^+^ HE cells, lower RUNX1 protein levels in HE cells, or a decrease in IAHC cells we analyzed *Tlr4*^*-/-*^ mouse embryos and their wildtype littermates by whole mount confocal microscopy (Fig. 1A-F). IAHC cells were identified by their rounded appearance and expression of RUNX1, CD31, and high levels of c-Kit, while HE cells were flatter, located within the endothelial layer, and expressed RUNX1, CD31, and very low or undetectable levels of c-Kit. We found that the median fluorescence intensity of RUNX1 protein was lower in HE cells in the DA of E10.5 *Tlr4*^*-/-*^ embryos compared to *Tlr4*^*+/+*^ littermates but the numbers of RUNX1^+^ HE cells were equivalent (Fig. 1A-C). The number of IAHC cells, on the other hand, was reduced by 42% in E10.5 *Tlr4*^*-/-*^ embryos, and IAHC cells expressed lower levels of RUNX1 protein (Fig. 1D,E). The reduction in IAHC cell number is consistent with observations in *tlr4bb* morphant zebrafish embryos, which had fewer myb:GFP^+^ kdrl:mCherry^+^ HSPCs in the DA (He et al., 2015).

**Figure 1.**
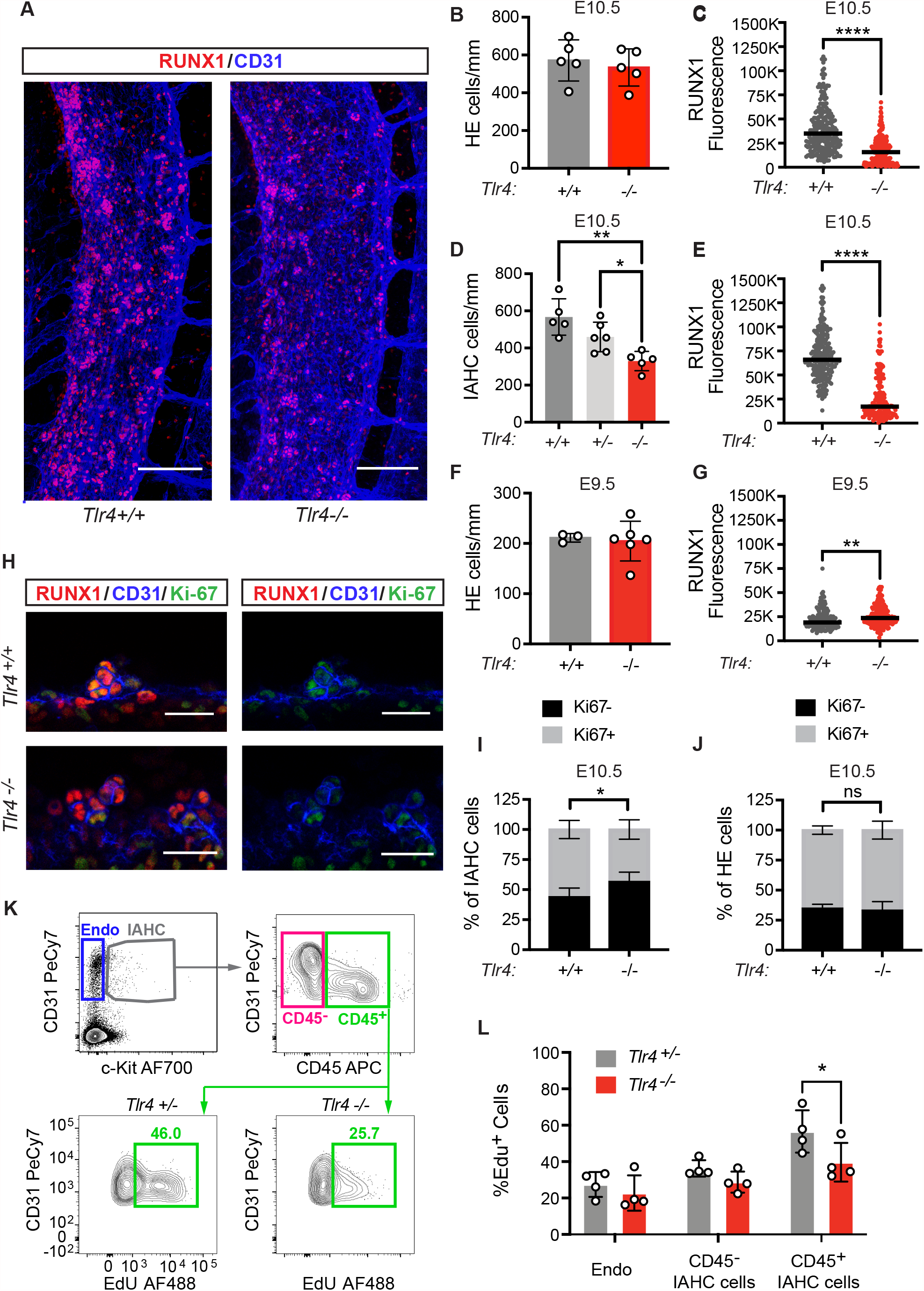
TLR4 signaling increases the number and proliferation of IAHC cells but does not regulate the number of HE cells. A) Representative images of confocal Z-projections (interval = 2µM) of the DA of E10.5 *Tlr4*^*+/+*^ (left) or *Tlr4*^*-/-*^ (right) embryos immunostained for CD31 (blue) and RUNX1 (red). Scale bar = 100µM. B) Quantification of HE cells (CD31^+^RUNX1^+^cKit^-^) per mm length of the DA in immunostained E10.5 embryos. Mean±SD; each dot represents one embryo; Student’s t-test, two-tailed; n=5 per genotype. The area counted centered on the intersection of the vitelline artery with the DA, and all cells within 2-3 somite pairs in both the rostral and caudal direction were counted. C) Corrected total cell fluorescence (CTCF) was calculated to measure RUNX1 intensity in HE cells described in (B). Integrated density values for each HE cell were measured using FIJI software and CTCF was calculated as [integrated density of cell - (area of selected cell x mean intensity of background)]. Bar represents the mean; Student’s t-test, two-tailed; n=248 for *Tlr4*^*+/+*^ and n=258 for *Tlr4*^*-/-*^ with cells measured from 3 embryos per genotype. D) Quantification of IAHC cells/mm (CD31^+^RUNX1^+^cKit^+^) in confocal images of E10.5 DAs from *Tlr4*^*+/+*^, *Tlr4*^*+/-*^ and *Tlr4*^*-/-*^ embryos. Mean±SD; one-way ANOVA and Tukey’s multiple comparison test, n=5-6 embryos per genotype. E) RUNX1 intensity determined by CTCF in IAHC cells in the DA as described in (D) for E10.5 *Tlr4*^*+/+*^ and *Tlr4*^*-/-*^ embryos. Bar represents the mean; Student’s t-test, two-tailed; n=223 for *Tlr4*^*+/+*^ and n=157 for *Tlr4*^*-/-*^ with approximately 50-60 cells measured from 3-4 embryos per genotype. F) Quantification of HE cells/mm at E9.5 in the DA of *Tlr4*^*+/+*^ and *Tlr4*^-/-^ embryos. Cells were counted in the region spanning the caudal boundary of the heart to the intersection with the umbilical artery. Mean±SD; Student’s t-test, two-tailed; n=3 and 6 for *Tlr4*^*+/+*^ and *Tlr4*^*-/-*^, respectively. G) CTCF values, indicating RUNX1 intensity, measured for HE cells in immunostained E9.5 *Tlr4*^*+/+*^ and *Tlr4*^-/-^ embryos. Bar represents the mean; Student’s t-test, two-tailed; n=141 for *Tlr4*^*+/+*^ and n=192 for *Tlr4*^*-/-*^ with approximately 40-50 cells measured from at least 3 embryos per genotype. H) Confocal Z-projections (2µm intervals, 4 intervals total/image) of IAHC cells in E10.5 DAs of *Tlr4*^*+/+*^ (top row) and *Tlr4*^-/-^ (bottom row) embryos immunostained with CD31 (blue), RUNX1 (red), and Ki67 (green). Scale bar = 25µm. I) Frequency of Ki67^+^ (gray) and Ki67^-^ (black) IAHC cells, calculated as [Ki67^+/-^ subset/total IAHC cells], observed in confocal images of E10.5 DAs of *Tlr4*^*+/+*^ and *Tlr4*^*-/-*^ embryos. Mean±SD; Two-way ANOVA, Sidak correction for multiple comparisons; n=6-7 per genotype. J) Frequency of Ki67^+^ (gray) and Ki67^-^ (black) HE cells, calculated as described in (I). K) Representative flow cytometry plots depicting gating for CD31^+^cKit^-^ endothelial cells (Endo), CD31^+^cKit^+^CD45^-^ IAHC cells, and CD31^+^cKit^+^CD45^+^ IAHC cells. EdU labeling of CD45^+^ IAHC cells (bottom row) are shown for *Tlr4*^*+/-*^ and *Tlr4*^*-/-*^ embryos. *Tlr4*^*+/-*^ embryos were used in lieu of *Tlr4*^*+/+*^ embryos in order to generate a sufficient number of cells for the analysis. The number of IAHC cells in *Tlr4*^*+/-*^ and *Tlr4*^*+/+*^ embryos were not significantly different, as shown in (D). Cells in plots were previously gated on FSC vs SSC, Live, lineage negative, CD41^lo/-^ (see Fig. S1). Representative of 3 independent experiments (4 litters). Littermates were pooled in each experiment. L) Frequency of EdU^+^CD31^+^cKit^-^ endothelial cells (Endo), CD31^+^cKit^+^CD45^-^ IAHC cells, and CD31^+^cKit^+^CD45^+^ IAHC cells determined by flow cytometry. Mean±SD; Two-way ANOVA, Sidak correction for multiple comparisons; n=4 pooled litters per genotype. In all panes, * p≤0.05, ** p≤0.01, *** p≤0.001, **** p≤0.0001,ns = not significant

The DA contains approximately 0-5 IAHC cells at E9.5 and ∼600 IAHC cells at E10.5 (Yokomizo and Dzierzak, 2010). Since the number of HE cells at a single snapshot in time is a function of the number that were originally specified minus the number that have undergone EHT, the optimal time for determining how many HE cells are specified is at E9.5, prior to the major egress of HE cells via EHT. The numbers of RUNX1^+^ HE cells were equivalent in E9.5 *Tlr4*^*+/+*^ and *Tlr4*^*-/-*^ embryos (Fig. 1F), and levels of RUNX1 in HE cells were slightly increased (Fig. 1G). Therefore, TLR4 deficiency does not affect the number of HE cells that are specified at E9.5, but it decreases RUNX1 levels in HE cells at E10.5.

The decreased number of IAHC cells in E10.5 *Tlr4*^*-/-*^ embryos could be caused by inefficient EHT, decreased proliferation, or increased apoptosis of IAHC cells. Inefficient EHT is unlikely as it would have resulted in an accumulation of HE cells in the DA in *Tlr4*^*-/-*^ embryos at E10.5, which we did not observe. *Tlr4*^*-/-*^ IAHC cells were, however, less proliferative, as reflected by a smaller fraction that were positive for the proliferation marker Ki-67 (Fig. 1H,I). Flow cytometric analysis also showed loss of TLR4 decreased proliferation, determined by EdU labeling, specifically of CD45^+^ IAHC cells, which at E10.5 lack pre-HSCs but are enriched for LMPs (Zhu et al., 2020) (Fig. 1K,L and Fig. S1). TLR4 deficiency did not affect the proliferation of endothelial cells, HE cells, or CD45^-^ IAHC cells, which at E10.5 include pre-HSCs (Fig. 1J,L). In summary, loss of TLR4 decreased RUNX1 protein levels in E10.5 HE and IAHC cells and also decreased the number and proliferation of CD45^+^ IAHC cells, but had no effect on the number of HE cells.

Both TLR4 and TLR2 are expressed in and regulate inflammatory signaling in endothelial cells (Salvador et al., 2016), therefore we determined whether TLR2 also regulates the number of IAHC cells. Loss of TLR2 reduced the number of IAHC cells in the DA to a similar extent as TLR4 deficiency (Fig. 2B). TLR2 signals exclusively through the MyD88 adaptor, while TLR4 signals through both the MyD88 and TRIF adaptors (Fig. 2A). To determine which signaling arm of TLR4 was important for regulating the number of IAHC cells we enumerated IAHC cells in both *Myd88*^*-/-*^ and TRIF-deficient (*Ticam1*^*-/-*^) embryos. Loss of MyD88, but not TRIF, decreased the number of IAHC cells (Fig. 2B). Therefore, signaling via cell surface TLR2 and TLR4 through the MyD88 adaptor, but not by endosomal TLR4 through the TRIF adaptor, regulates the number of IAHC cells.

**Figure 2.**
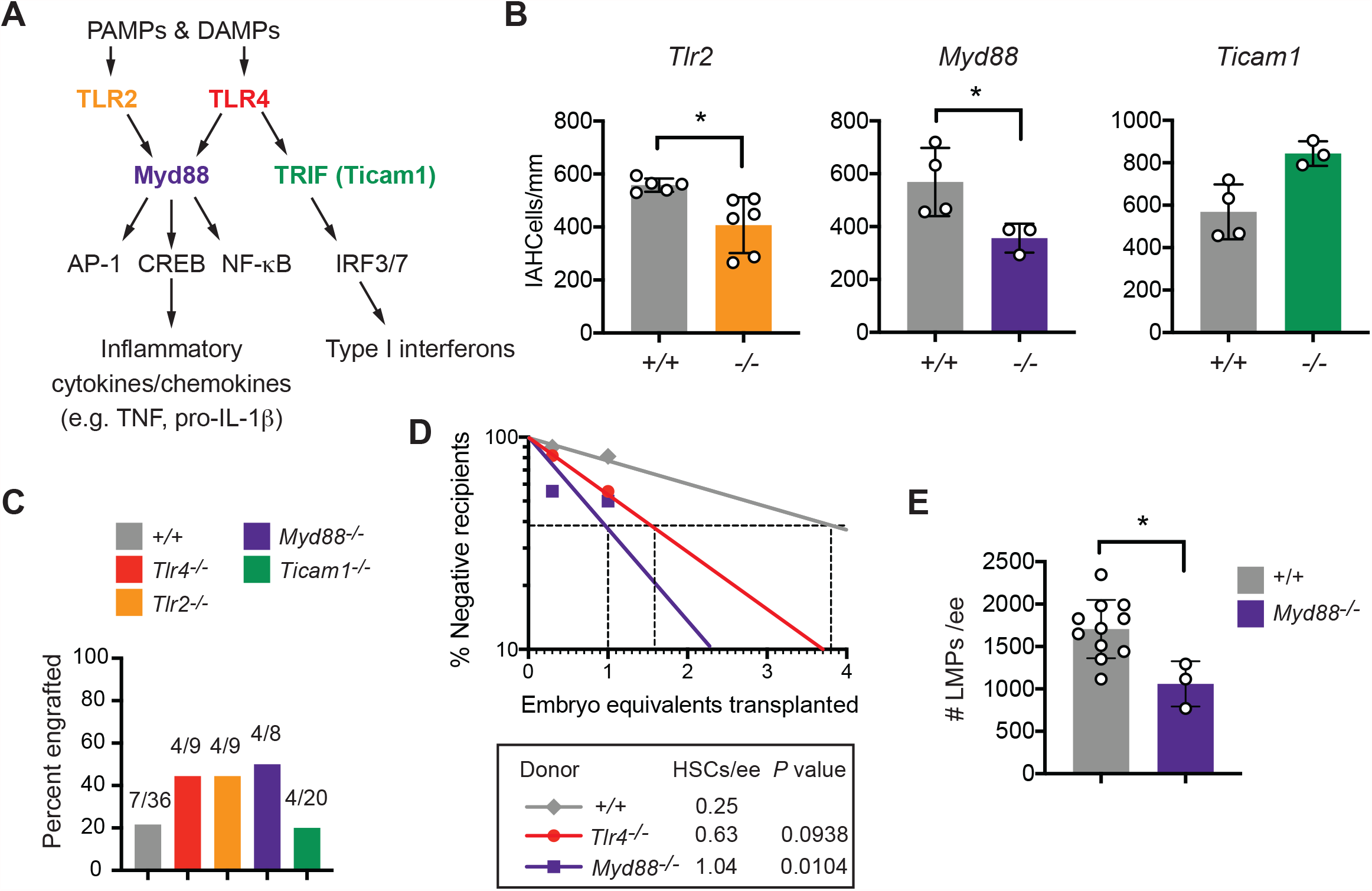
MyD88-dependent TLR signaling increases the number of LMPs in the AGM region but limits the number of HSCs. A) TLR signaling pathway highlighting cell surface receptors TLR2 and TLR4, their adaptors MyD88 and TRIF, and transcription factor effectors. B) Quantification of CD31^+^RUNX1^+^cKit^+^ IAHC cells/mm in the DA of E10.5 embryos from confocal images. Student’s t-test, two-tailed, n=3-6 per genotype. C) Frequency of recipient mice (CD45.1^+^) exhibiting multi-lineage reconstitution following transplantation of 1 ee of AGM regions from E11.5 embryos (CD45.2^+^). Multi-lineage reconstitution was assessed as >1% donor myeloid (Mac1^+^, Mac1^+^Gr1^+^), T-cells (CD3^+^), and B-cells (CD19^+^) in the peripheral blood 16 weeks post-transplantation (see Fig. S2 for details of flow cytometric analysis). D) Quantification of HSCs in E11.5 wild type (+/+), *Tlr4*^*-/-*^, and *MyD88*^*-/-*^ embryos by limiting dilution transplantation of 0.3 and 1.0 ee cells from the AGM region, umbilical and vitelline arteries (CD45.2^+^) into CD45.1^+^ adult recipients. Representative of at least 8 recipients per group from 7 transplants. The number of HSCs was calculated using ELDA software (Hu and Smyth, 2009). E) Frequency of LMPs in E10.5 +/+ and *Myd88*^*-/-*^ embryos (n=11, n=3, respectively) determined by plating embryos in limiting dilution on OP9 and OP9-DL1 stromal cells. +/+ embryos include offspring from intercrosses of mice heterozygous for mutations in other inflammatory mediators.

### MyD88-dependent TLR signaling positively regulates the number of committed lymphoid progenitors but restrains the number of adult repopulating HSCs

To determine whether loss of TLR signaling also decreased the number of adult-repopulating HSCs capable of multilineage engraftment, we transplanted 1 embryo equivalent (ee) of aorta-gonad-mesonephros (AGM) regions from E11.5 wild type, *Tlr4*^*-/-*^, *Tlr2*^*-/-*^, *Myd88*^*-/-*^, and *Ticam1*^*-/-*^ embryos into irradiated adult recipient mice, and measured contribution to peripheral blood and bone marrow 16 weeks post-transplant (Fig. S2). One E11.5 wild type AGM region contains ∼1 adult repopulating HSC, and therefore, 1ee is a limiting dose (Kumaravelu et al., 2002). Despite the decrease in the number of IAHC cells, loss of TLR4, TLR2, and MyD88 increased the proportion of mice with multi-lineage engraftment (Fig. 2C), indicating that adult-repopulating HSCs were not depleted in the AGM regions of TLR4, TLR2, or MyD88-deficient mouse embryos but instead appeared to be increased. Loss of *Ticam1*, on the other hand, which did not significantly change IAHC cell numbers, did not affect multi-lineage engraftment (Fig. 2B,C).

To quantify the numbers of multi-lineage adult repopulating HSCs in embryos with mutations affecting MyD88-dependent TLR signaling, we transplanted a cohort of mice with limiting doses (0.3 ee and 1.0 ee) of dissociated cells from the AGM regions of wild type, *Tlr4*^*-/-*^ and *Myd88*^*-/-*^ embryos. There was a trend towards increased numbers of HSCs in the AGM regions of E11.5 *Tlr4*^*-/-*^ embryos and a significant 4.2-fold increase in the number of multi-lineage repopulating HSCs in *Myd88*^*-/-*^ embryos (Fig. 2D).

LMPs with the potential to produce B and T cells in culture are >10-fold more abundant in AGM regions of E10.5 embryos compared to HSCs and pre-HSCs, and are enriched in the CD45^+^ population (Li et al., 2014; Rybtsov et al., 2016; Zhu et al., 2020). To determine whether the number of LMPs was altered by mutations affecting MyD88-dependent signaling we performed a limiting dilution analysis of cells from the AGM regions of wild type and *Myd88*^*-/-*^ embryos by plating them into wells containing OP9 stromal cells to enumerate progenitors with B cell potential, and on OP9 cells expressing the Notch ligand delta-like 1 (OP9-DLL1) to measure the number of T cell progenitors (Li et al., 2014; Schmitt and Zuniga-Pflucker, 2002). *Myd88*^*-/-*^ embryos contained significantly fewer progenitors capable of producing B or T cells compared to wild type embryos (Fig. 2E). Therefore, MyD88-dependent TLR signaling increases the generation of LMPs at the expense of long-term multilineage-repopulating HSCs.

### TLR4 signaling in endothelial cells and/or IAHCs, but not in primitive innate immune myeloid cells, regulates IAHC cell numbers

Professional innate immune myeloid cells are the primary producers of inflammatory cytokines and chemokines and are proposed to be an important cellular source of the inflammatory molecules promoting the generation of HSPCs in the DA. Depletion of macrophages and granulocytes in zebrafish embryos using morpholinos targeting *spi1a* +*spi1b*, of macrophages alone with morpholinos against *irf8*, or of neutrophils using morpholinos to *cebp1* decreased *runx1* expression in the DA and the number of myb:GFP^+^ kdrl:mCherry^+^ HSPCs (Espin-Palazon et al., 2014; He et al., 2015; Li et al., 2014). Depletion of macrophages in explanted AGM regions from mouse embryos by clodronate also decreased HSPCs (Mariani et al., 2019). Vital imaging documented macrophages intimately contacting IAHC cells, and macrophages were occasionally observed to extravasate through endothelial cells to reach the IAHCs (Mariani et al., 2019; Travnickova et al., 2015).

We determined whether TLR4 signaling in yolk sac-derived myeloid cells regulates the number of IAHC cells in the DA by deleting *Tlr4* floxed alleles (*Tlr4*^*f/f*^) in EMPs using Csf1r-Cre (Gomez Perdiguero et al., 2015; Schulz et al., 2012). Csf1r-Cre should result in TLR4 loss in both yolk sac-derived macrophages and granulocytes, although since granulocytes are a minor population in the mouse embryo, macrophages are the more likely source of inflammatory cytokines (Mariani et al., 2019). We confirmed that cell surface TLR4 was depleted on macrophages following Csf1r-Cre deletion (Fig. 3A, Fig. S3A), and that macrophages had an impaired response to LPS stimulation, as evidenced by lower TNF production (Fig. 3B). Deletion of TLR4 by Csf1r-Cre had no impact on the number of IAHC cells (Fig. 3C, Fig. S3B), indicating that the cytokines produced by primitive myeloid cells in response to TLR4 signaling do not regulate the number of IAHCs. RUNX1 levels were significantly lower in IAHCs, but whether this was due to Csf1r-Cre mediated deletion in EMPs, or in differentiating IAHC cells could not be determined (Fig. 3D). These results do not, however, exclude a role for primitive myeloid cells in regulating IAHC cell numbers, as other signaling pathways that regulate inflammatory cytokine production or other processes mediated by primitive myeloid cells may be responsible for their effect on hematopoietic ontogeny in the DA.

**Figure 3.**
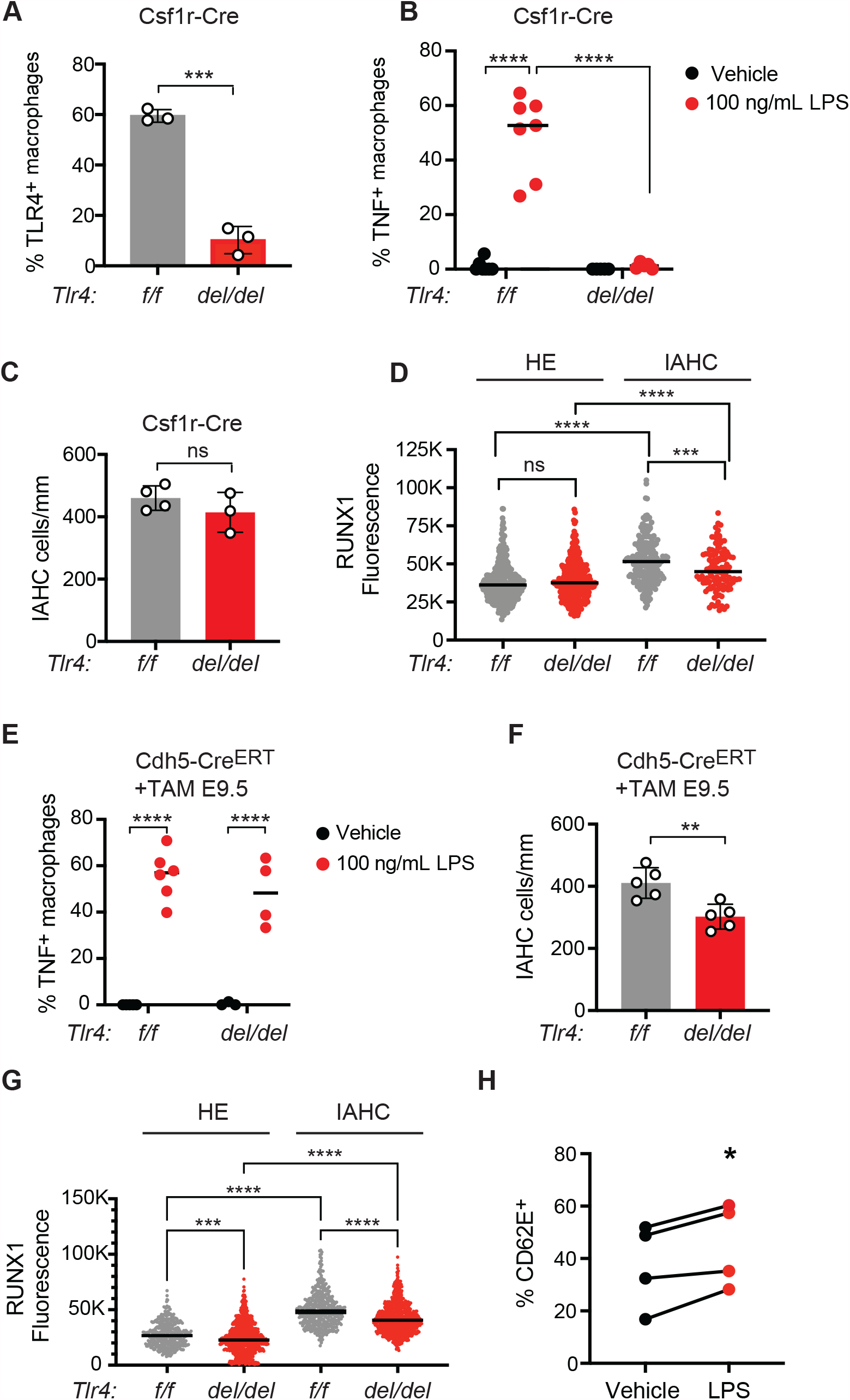
TLR4 signaling is required in endothelial and/or IAHC cells to promote IAHC cell numbers. A) Percent TLR4^+^ macrophages (CD45^+^Mac-1^+^F4/80^+^) measured by flow cytometry in E10.5 *Tlr4*^*f/f*^ (*f/f*) and *Tlr4*^*f/f*^;Csf1r-Cre embryos (*del/del*). Mean±SD; 3 independent experiments with pooled littermates; Student’s t-test, two-tailed (see Fig. S3A for flow plots.) B) Frequency of TNF^+^ macrophages in E10.5 *Tlr4*^*f/f*^ and *Tlr4*^*f/f*^;Csf1r-Cre embryos treated *ex vivo* with vehicle or LPS, determined by flow cytometry (see Fig. S3B for flow plots). Bar represents the mean; Two-way ANOVA with Sidak’s correction for multiple comparisons; n=5-7. C) Quantification of CD31^+^RUNX1^+^c-Kit^+^ IAHC cells in the DA of E10.5 *Tlr4*^*f/f*^ and *Tlr4*^*f/f*^;Csf1r-Cre embryos. Mean±SD; Student’s t-test, two-tailed; n=3. D) CTCF measuring RUNX1 intensity in HE and IAHC cells in E10.5 *Tlr4*^*f/f*^ and *Tlr4*^*f/f*^;Csf1r-Cre embryos. Two-way ANOVA with Sidak’s correction for multiple comparisons; HE cells: n=378 and 333, IAHC cells n=209 and 119 for *Tlr4*^*f/f*^ and *Tlr4*^*f/f*^;Csf1r-Cre, respectively. E) Frequency of TNF^+^ macrophages in vehicle and LPS-stimulated explanted E10.5 *Tlr4*^*f/f*^ and *Tlr4*^*f/f*^;Cdh5-Cre^ERT^ embryos. Dams were injected with tamoxifen (TAM) at E9.5. Bars represent the mean; Two-way ANOVA with Sidak’s correction for multiple comparisons; n=3-6. F) Quantification of IAHC cells (CD31^+^RUNX1^+^c-Kit^+^) in E10.5 *Tlr4*^*f/f*^ and *Tlr4*^*f/f*^;Cdh5-Cre^ERT^ embryos. Mean±SD; Student’s t-test, two-tailed; n=5. G) RUNX1 intensity measured as CTCF in HE and IAHC cells in E10.5 *Tlr4*^*f/f*^ and *Tlr4*^*f/f*^;Cdh5-Cre^ERT^ embryos. Two-way ANOVA with Sidak’s correction for multiple comparisons; HE: n=389 and 739, IAHC n=454 and 789 cells for *Tlr4*^*f/f*^ and *Tlr4*^*f/f*^;Cdh5-Cre^ERT^ respectively. H) Frequency of CD62E^+^ endothelial cells following 6 hour stimulation of FACS-purified endothelial cells with LPS (100 ng/mL). Represents 4 independent experiments. Paired Student’s t-test, two-tailed.

We next assessed whether TLR4 signaling in endothelial or IAHC cells was necessary for generating normal numbers of IAHC cells by deleting *Tlr4*^*f/f*^ alleles with an endothelial cell-specific, tamoxifen-regulated Cre (Cdh5-Cre^ERT^) (Sorensen et al., 2009a). We timed the tamoxifen injections in pregnant dams to preferentially delete TLR4 in arterial endothelial cells and IAHCs but not in EMPs. EMPs differentiate from HE cells in the yolk sac vascular plexus between E8.5-E9.5 (Frame et al., 2015), just prior to the major wave of EHT in arterial endothelium that produces IAHC cells (Tober et al., 2013; Yokomizo and Dzierzak, 2010). We previously performed a temporal analysis of Cdh5-Cre^ERT^ activity following tamoxifen injection and demonstrated that deletion in EMPs could be mitigated by injecting tamoxifen at E9.0 or later in development (Tober et al., 2013). Therefore, to delete TLR4 in arterial endothelial cells while sparing EMPs we injected tamoxifen into pregnant dams at E9.5 and analyzed HE and IAHC cells one day later in *Tlr4*^*f/f*^ and *Tlr4*^*f/f*^; Cdh5-Cre^ERT^ embryos. We confirmed that TNF production was unchanged in macrophages from *Tlr4*^*f/f*^; Cdh5-Cre^ERT^ embryos following stimulation with LPS at E10.5, confirming that deletion of *Tlr4* using a carefully timed activation of Cdh5-Cre^ERT^ has minimal impact on EMPs or their progeny (Fig. 3E). The number of IAHC cells at E10.5 was significantly decreased following deletion of *Tlr4*^*f/f*^ alleles at E9.5 by Cdh5-Cre^ERT^, as were RUNX1 levels in HE and IAHC cells (Fig. 3F,G). We conclude that TLR4 is required in arterial endothelial cells and/or IAHC cells to generate normal numbers of IAHC cells.

To confirm that TLR4 signaling can be directly activated in embryonic endothelial cells we purified CD31^hi^VEC^+^ESAM^+^CD44^-^CD45^-^Ter119^-^Mac1^-^CD41^-^ endothelial cells, stimulated them *ex vivo* with LPS and monitored expression of the integrin CD62E (E-selectin, encoded by *Sele*) which is a direct NF-κB target gene (Collins et al., 1995). CD62E levels increased in purified endothelial cells in response to LPS compared to unstimulated endothelial cells (Fig. 3H), indicating that embryonic endothelial cells respond to TLR4 signals. We were unable to accurately detect TLR4 by flow cytometry on the surface of endothelial cells. However, since signaling through TLR4 on the cell surface requires MyD88, and the genetic data indicate that TLR signaling through MyD88 but not TRIF adaptors regulates IAHC numbers, we conclude that the relevant signaling is occurring through extracellular TLR4 signaling.

In summary, the number of cells in the IAHCs, and the number of LMPs are positively regulated by MyD88-dependent TLR signaling (Fig. 4). The number of specified HE cells, on the other hand, is not. MyD88-dependent TLR signaling is required in endothelial cells or in the IAHC cells that differentiate from HE cells to regulate IAHC cell numbers. The selective decrease in LMPs in the absence of MyD88-dependent signaling inversely correlates with an increase in adult-repopulating HSCs, suggesting that MyD88-dependent TLR signaling regulates the balance between stem cells and progenitors with different potentials (Fig. 4). Previous single cell RNA sequencing demonstrated that LMPs express higher levels of proliferation genes, such as *Myc*, compared to pre-HSCs/HSCs (Zhu et al., 2020). Loss of TLR4 signaling and decreased proliferation may differentially affect both committed progenitors and pre-HSCs/HSCs. TLR4 signaling in adult HSCs increases proliferation but reduces fitness (Takizawa et al., 2017), therefore decreased proliferation of pre-HSCs and HSCs in MyD88-deficient embryos may contribute to the selective increase in functional HSCs.

**Figure 4.**
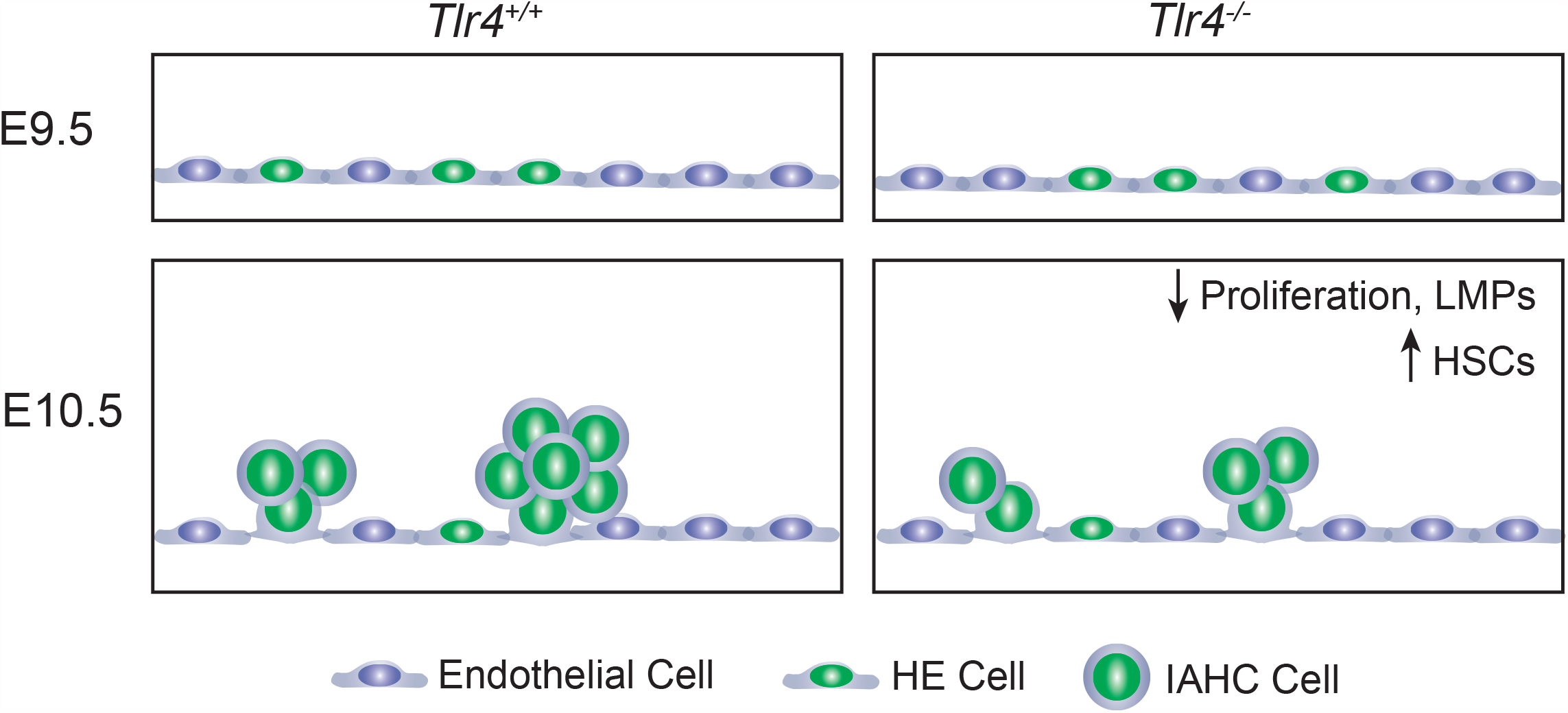
TLR4 signaling increases IAHC cells via proliferation, increases RUNX1 levels and promotes the formation of LMPs while repressing HSC numbers.

Haploinsufficiency for RUNX1 decreases the number of IAHC cells and results in the precocious emergence of HSCs in the embryo (Cai et al., 2000). However, Runx1 haploinsufficiency also decreases the number of HE cells (Zhu et al., 2020), whereas loss of MyD88-dependent TLR signaling does not. HE cells differentiate from a precursor population called pre-HE, and the efficiency at which pre-HE differentiates into HE is regulated by RUNX1 dosage (Zhu et al., 2020). We believe the reason loss of MyD88-dependent TLR signaling does not reduce the number of HE cells is that RUNX1 levels are normal in mutant embryos at E9.5 when specification occurs. The subsequent decrease in RUNX1 levels in HE and IAHC cells at E10.5 may contribute to the lower number and proliferation of IAHC cells, and reflect a later requirement for RUNX1. RUNX1 promotes the G_1_ to S phase transition in several contexts (Cai et al., 2011; Nimmo et al., 2005; Osorio et al., 2008), and loss of RUNX1 decreases proliferation, consistent with these observations.

Our data are consistent with those from zebrafish embryos which demonstrated lower levels of *runx1* expression and decreased numbers of HSPCs upon loss of MyD88-dependent TLR signaling (He et al., 2015), but they add an extra layer of complexity. They also provide a cautionary note for investigators studying HSC ontogeny--that the level of RUNX1 expression, and the numbers of IAHC cells and progenitors do not always correlate with the number of HSCs.

## Methods

### Mouse lines and embryo production

*Tlr4*^*+/-*^ (B6(Cg)-*Tlr4*^*tm1*.*2Karp*^/J), *Tlr4*^*f/f*^ (B6(Cg)-*Tlr4*^*tm1*.*1Karp*^/J), *Csf1r*^*Cre*^ (C57BL/6-Tg(Csf1r-cre)1Mnz/J), *Ticam1*^*+/-*^ (C57BL/6J-*Ticam1*^*Lps2*^/J), and B6.SJL-*Ptprc*^*a*^ *Pepc*^*b*^*/*BoyJ (CD45.1) mice were purchased from JAX. Tg(Cdh5-cre/ERT2)1Rha mice (Sorensen et al., 2009b) were a gift from Ralf Adamson. *Myd88*^*+/-*^ mice were a gift from Edward Behrens.

### Animal husbandry

The morning post mating is considered embryonic day (E) 0.5. E9.5-E11.5 embryos were staged at the time of harvest by counting somites. Embryos that showed abnormal development were discarded. Mice were handled according to protocols approved by the University of Pennsylvania’s Institutional Animal Care and Use Committee and housed in a specific-pathogen-free facility.

### Cell sorting and flow cytometry

Antibodies used can be found in Supplemental Table 1. All antibodies were used at a concentration of 1:200. DAPI, LIVE/DEAD Fixable Aqua (ThermoFisher L34957), or Fixable Viability Stain 780 (BD Biosciences 565388) were used to exclude dead cells. We used the Cytofix/Cytoperm Kit (BD Biosciences) for intracellular flow cytometry to detect TNF in cells from embryos that were treated *ex vivo* with 100ng/mL LPS or vehicle for 4 hours.

Yolk sacs were removed from embryos, and vitelline vessels were retained with the embryonic portion. The head, cardiac and pulmonary regions, liver, digestive tube, tail and limb buds were removed. The remaining portion containing the aorta-gonad-mesonephros (AGM) region, portions of somite, umbilical and vitelline vessels were collected. Tissues were dissected in phosphate buffered saline (PBS)/10% Fetal Bovine Serum (FBS) and Penicillin/Streptomycin (Sigma), followed by dissociation in 0.125% Collagenase (Sigma) for 1 hour. Tissues were washed and filtered through a 40-micron filter and resuspended in antibody solution. Cells were sorted on a BD Influx equipped with a 100-micron nozzle and run at a pressure of 17 psi with flow rates less than 4000 events/second and collected in PBS/20% FBS/25mM HEPES. Flow cytometry was performed on an LSR II (BD) and data were analyzed using FlowJo software (v10.7, BD).

### T and B cell progenitor assays

OP9 and OP9-DL1 stromal cells were maintained in αMEM containing 20% FBS and antibiotics and plated into 96-well plates at a concentration of 5500 cells/well 1 day before dilutions of the dissociated embryonic tissues were added. Medium was supplemented with purified recombinant 5ng/ml Flt3L and 10ng/ml IL-7 (OP9) or 5ng/ml Flt3L and 1ng/ml IL-7 (OP9-DL1) (Peprotech). Cells were harvested and analyzed by flow cytometry 12 days after culture. B cells were identified as CD19^+^ B220^+^, T cells as CD25^+^ Thy1.1^+^. The progenitor frequency was calculated using the method of maximum likelihood applied to Poisson distribution with L-Calc software (StemCell Technologies). OP9s were purchased from ATCC, and OP9-DL1s were obtained from Juan Carlos Zún̋iga-Pflücker.

### EdU labeling

Yolk sacs were removed and intact embryos were placed in αMEM containing 20%FBS and antibiotics. Embryos were incubated in a 24-well plate at 37°C for 2 hours in EdU (5-ethynyl-2’
s-deoxyuridine, 100uM). Immediately following incubation with EdU, embryos were dissociated using 0.125% collagenase (Sigma) for 30 minutes and stained for flow cytometry. EdU detection was performed using the Click-iT Plus EdU Flow Cytometry Assay kit (Thermo Fisher, C10632).

### Transplant analyses

B6.SJL-*Ptprc*^*a*^*Pepc*^*b*^/BoyJ (CD45.1) mice were subjected to a split dose of 900 cGy, 3 hours apart. Each recipient received 1.0 or 0.3 embryo equivalents (ee) of E11.5 AGM regions (CD45.2), vitelline and umbilical arteries with 2.5×10^5^ carrier spleen cells by retro-oribital injection. We assessed donor (CD45.2) engraftment in peripheral blood at weeks 4, 8, 12 and 16, and in bone marrow at week 16 post transplantation. HSC frequencies were determined by extreme limiting dilution analysis (ELDA) (Hu and Smyth, 2009).

### Confocal Microscopy

Embryos were prepared as previously described (Yokomizo et al., 2012). A Zeiss LSM 880 AxioObserver inverted confocal microscope with ZEN 2.3 SP1 software was used to acquire Z-projections and single optical projections. Images were processed using Fiji software (Schindelin et al., 2012). The following primary antibodies were used; rat anti-mouse CD31 (AB_396660, 1:500), rat anti-mouse CD117 (AB_467434, 1:250), rabbit anti-human/mouse RUNX (AB_2049267, 1:500), and rabbit anti-mouse Ki67 (AB_443209, 1:500). Secondary antibodies used were goat anti-rat Alexa Fluor 647 (AB_2864291), goat-anti rat Alexa Fluor 555 (AB_150158), goat anti-rabbit Alexa Fluor 488 (AB_2630356), and goat anti-rabbit Alexa Fluor 647 (AB_2535813). All secondaries were used at a concentration of 1:1000.

### Statistics

Significance for multiple comparisons was determined using ordinary one-way ANOVA with Tukey’s test for multiple comparisons or two-way ANOVA with Sidak’s correction for multiple comparisons. Two-tailed student’s t-test was used for pairwise comparisons. All statistics were performed using Prism software (v. 9.0.0). For all tests, * = p<0.05, ** = p<0.01, *** = p<0.001, **** = p<0.0001.

## Acknowledgments

This work was supported by National Institutes of Health grants R01HL091724. L.B. is supported by T32HL007439 and M.M. by T32DK007780.

## Authorship

### Contribution

L.B, M.M., and Y.L. conducted the experiments. N.A.S. directed the project and wrote the manuscript.

## Figure Legends

**Supplemental Figure 1. Proliferation is decreased in CD45**^**+**^ **IAHC cells**.

A) Representative flow plots depicting gating strategy of endothelial cells (Endo), CD45^-^ and CD45^+^ IAHC cells and the percent of EdU^+^ cells in each population in *Tlr4*^*+/-*^ and *Tlr4*^*-/-*^ embryos. Numbers on the x and y-axes are indicated on the first plot on the left, and unless changed are not depicted on plots to the right of the preceding plot.

**Supplemental Figure 2. Loss of TLR4 does not affect adult-repopulating HSCs**.

A,B) Representative flow plots depicting gating strategy of myeloid, B and T cells from donor cells and the frequency of donor cells comprising each lineage. Numbers on the x and y-axes are indicated on the first plot on the left, and unless changed are not depicted on plots to the right of the preceding plot.

**Supplemental Figure 3. Csf1r-Cre effectively removes cell surface TLR4 and TLR4 signaling in embryonic macrophages**.

A) Representative flow plots depicting gating strategy to examine surface TLR4 on embryonic macrophages described in Fig. 3A. Numbers on the x and y-axes are indicated on the first plot on the left, and unless changed are not depicted on plots to the right of the preceding plot.

B) Representative flow plots depicting gating strategy to examine TNF production in macrophages in response to LPS stimulation.

